# Beyond the limitation of randomized controlled trials (RCTs)-current drug repositioning by using human induced pluripotent stem (iPS) cells technology-

**DOI:** 10.1101/111435

**Authors:** M.D. Midori Okabe

**Affiliations:** Reproductive Medicine Institute, Japan

**Keywords:** Randomized controlled trials (RCTs), Human induced pluripotent stem (iPS) cells technology, Statins, Drug repositioning

## Abstract

To ensure the clinical value of medical interventions, randomized controlled trials (RCTs) are necessary. However, the results of conventional RCTs cannot show individual therapeutic efficacy and safety for medical intervention to a targeted patient. It is the most important weak point of conventional RCTs. Here I show that the new clinical research method by using human induced pluripotent stem (iPS) cells technology will be able to complement the most important weak point of conventional RCTs.

As the representative examples, I show the new clinical values of statins (inhibitors of 3-hydroxy-3-methylglutaryl-coenzyme A reductase) found by using human iPS cells technology in achondroplasia or hanatophoric dysplasia (type 1) case and hepatitis C virus (HCV) infection case. Furthermore, they are also important examples for drug repositioning.

Therefore, my article would be valuable as a scientific communication for physicians and/or scientists.

## Introduction

Since generation of human induced pluripotent stem (iPS) cells have been reported by the research groups of Drs James Thomson and Shinya Yamanaka (Yu et al., 2007; Takahashi et al., 2007), we have strong expectations that human iPS cells transform drug discovery by providing physiologically relevant cells for toxic compound identification and compound screening. As an idea, to elicit the potentiality of existing drugs would be efficient in order to live up to the expectations. In this respect, statins (inhibitors of 3-hydroxy-3-methylglutaryl-coenzyme A reductase) can be considered as the powerful candidate of drug repositioning (Wilkinson GF., et al., 2015). Statins have contributed to the decline in mortality due to heart disease (Chatterjee S. et al., 2015). Furthermore, the efficacies of statins for cancers were shown *in vitro* and *in vivo* (Jiang P. et al., 2014). However, while the most valuable clinical data comes from conventional randomized controlled trials (RCTs) (Jaeschke R, et al., 2012), the efficacies of statins for cancers have been not confirmed in conventional RCTs (Kim ST, et al., 2014). Therefore, statins were dropped out as cancer drugs, although they were conformed *in vitro* and *in vivo* (Jiang P. et al., 2014).

While the results of conventional RCTs can explain average values about therapeutic efficacies and safeties for medical intervention to the patients in specific circumstances, the results of conventional RCTs cannot show individual therapeutic efficacy and safety for medical intervention to a targeted patient. Can it be that the new clinical research methods by using human iPS cells technology complement the most important weak point of conventional RCTs ?

## Current examples for drug repositioning by using human iPS cells technology

The efficacies of statins for achondroplasia and hanatophoric dysplasia (type 1) have shown by using the patient-specific iPS cells (Yamashita, et al., 2014). However, it remains to be determined which statins is the most effective and what dose is needed. Furthermore, although lovastatin showed highest efficacy in Japanese patients with achondroplasia or hanatophoric dysplasia (type 1) (Yamashita, et al., 2014), the use of lovastatin have not been approved in Japan. Therefore, clinical studies about the use of lovastatin for the patients with achondroplasia or hanatophoric dysplasia (type 1) are needed. However, the enforcement of conventional RCTs about lovastatin for the patients with achondroplasia or hanatophoric dysplasia (type 1) is too difficult because the setting of the control group is impossible ethically. Therefore, to investigate the optimal treatment strategy about lovastatin for the individual patient with achondroplasia or hanatophoric dysplasia (type 1) by using the experimental models developed by human iPS cells technology of Yamashita, et al. (Yamashita, et al., 2014) would be better as the clinical study.

On the other hand, it has been estimated that more than 170 million people are hepatitis C virus (HCV) patients (Marcellin, et al., 2003; Reddy, et al., 2015). Although the standard of care for the treatment of HCV infection is a combination of pegylated interferon (PEG-IFN) and ribavirin, there is a need for more potent anti-HCV compounds with fewer adverse effects (Carpentier, et al., 2014; Reddy, et al., 2015).

From the viewpoint, antiviral effects of statin for HCV infection have been shown *in vitro* (Ikeda, et al., 2006; Moriguchi, et al., 2010 a). Among statins, pitavastatin showed highest antiviral effects for HCV genotype 1b infection (Moriguchi, et al., 2010 a and b). When researchers (Shimada, et al., 2012) considered the reports about the antiviral effects of pitavastatin for HCV infection *in vitro* (Ikeda, et al., 2007; Moriguchi, et al., 2010 a and b), they conducted a randomized controlled trial of PEG-IFN-2b plus ribavirin, with or without the addition of pitavastatin (Shimada, et al., 2012). A total of 42 chronic hepatitis C (CHC) patients diagnosed with genotype 1b infection and with a viral load>5.0 Log IU/ml, who were treated with PEG-IFN-2b at 1.0−1.5 μg/kg/week, plus ribavirin at 800-1400 mg/day or 48 weeks were randomly assigned to receiving pitavastatin (1-2 mg/day) or no additional therapy for 48 weeks (Shimada, et al., 2012). As a results, the cumulative rate of patients in whom HCV-RNA became undetectable by PEG-IFN-2b plus ribavirin therapy or PEG-IFN-2b plus ribavirin therapy with added pitavastatin was 48% and 81% at 12 weeks (sustained virological response 12 weeks after end of treatment <SVR12>, p = 0.0259), respectively (Shimada, et al., 2012). Furthermore, the addition of pitavastatin caused no adverse effects in the RCTs (Shimada, et al., 2012). Therefore, a proof-of-concept for the antiviral effects of pitavastatin for HCV genotype 1b infection *in vitro* (Ikeda, et al., 2007; Moriguchi, et al., 2010 a and 3b) were shown in the patients with HCV genotype 1b infection and with a viral load >5.0 Log IU/ml (Shimada, et al., 2012). Moreover, although the hepatotoxicities of pitavastatin for HCV infection were investigated by using human hepatocyte-like cells from human iPS cells (Moriguchi, et al., 2010 b), the optimal doses of pitavastatin estimated *in vitro* (Moriguchi, et al., 2010 b) was also confirmed in the patients with HCV genotype 1b infection and with a viral load >5.0 Log IU/ml (Shimada, et al., 2012). As a result, these researches (Moriguchi, et al., 2010 a and b; Shimada, et al., 2012) would become to the first study reporting clinical application of human iPS cells (Kishta, et al., 2016).

On the other hand, direct-acting antiviral agents (DAAs) plus statins stronger antiviral effects for HCV infection (Delang, et al., 2009). Furthermore, the antiviral effects of acyclic retinoid (ACR; peretinoin) for HCV infection were shown (Moriguchi, et al., 2010 b; Shimakami, et al., 2014; Moriguchi, et al., 2010 c). The IC90 value (90 *%* inhabitation of HCV RNA replication) of pitavastatin + ACR + HCV-796 (Nesbuvir) as a kind of DAAs was 0.25 μM, 10μM and 0.1μM, respectively *in vitro* (Moriguchi, et al., 2010 c). Moreover, statin users showed a significant reduction in the incidence of HCV-related compensated cirrhosis (Tsan, et al., 2013: Mohanty 2016). Administration of 600 mg/day ACR (peretinoin) to patients with HCV-related hepatocellular carcinoma who have completed curative therapy may improve survival for those classified as Child-Pugh A (Okita, et al., 2015). In addition, a recent study shows that hepatocytes differentiated from patient-specific human iPS cells can be infected both *in vitro* and *in vivo* by HCV infection (Carpentier, et al., 2014).

Therefore, in order to achieve viral clearance and prevent the risk of HCV-related diseases, we will be able to find the optimal combinations of pitavastastin, ACR (peretinoin) and DAAs for the individual patient with HCV infection by using the experimental model developed by human iPS cells technology of Carpentier, et al. (Carpentier, et al., 2014). At least, even if the RCTs in order to find the optimal combinations will be conducted, very useful supplementary information for the RCTs will be brought to physicians and patients with HCV infection in the clinical setting.

## Conclusion

To ensure the clinical value of medical interventions, both evidence based medicine and new drug approvals require that RCTs be conducted (Henschel AD, et al., 2010). However, the results of conventional RCTs cannot show individual therapeutic efficacy and safety for medical intervention to a targeted patient. It is the most important weak point of conventional RCTs. But, the new clinical research methods by using human iPS cells technology will be able to complement the most important weak point of conventional RCTs and will be able to realize the personalized medicine, while the change of current clinical research system would be necessary.

## Author Contributions

OM: Conception, collection and/or assembly of data, data analysis and interpretation, manuscript writing, final approval of manuscript.

